# The Role of Sex in Atherosclerotic Inflammation and Lipid-Handling Dysfunction

**DOI:** 10.1101/2024.08.22.608101

**Authors:** Phydena Z Liu, Lauren N Zhang, Alyssa J Matz

**Author notes:** Equal Contribution (Phydena Z Liu, Lauren N Zhang).

## Abstract

Residual risk of cardiovascular events remains despite treatments that effectively lower cholesterol levels and hypertension, suggesting that there must be more variables to consider in atherosclerosis treatment. Several studies have suggested sex^1,2,3^ and inflammation^4,5^ as important variables. However, a cross-cohort analysis of sex and risk factors like inflammation and lipid-handling dysfunction is needed to strengthen their connection to atherosclerosis. By using blood transcriptomic profiles of 391 male and female participants, this study revealed that inflammation and lipid-handling dysfunction have sex-specific roles in atherosclerosis. Transcriptomics of 391 human blood samples with varying degrees of atherosclerosis were used to identify sex-specific changes in immune response and lipid-handling in circulating blood cells. Preliminary analyses of both FPKM and normalized counts datasets showed that inflammatory pathway activation and enrichment increased as atherosclerotic disease severity increased across all sexes. Analysis of sex-specific differentially expressed genes (DEGs) using IPA’s Canonical Pathways showed that severely impacted females had more enriched inflammatory pathways than severely impacted males. Further cross-cohort analysis of sex-specific inflammation and lipid-handling dysfunction was performed using AtheroSpectrum, a single-sample macrophage annotation tool. AtheroSpectrum confirmed that inflammation was more critical to female atherogenesis and revealed that lipid-handling dysfunction was more critical to male atherogenesis. Our study underscored the importance of inflammation and sex as variables to consider in atherosclerosis treatment, suggesting that treatment should target inflammation and consider sex. Our findings may be used for generating a model to predict atherosclerosis risk based on key DEGs, pathways, sex, and other clinical parameters when available.

## Introduction

Atherosclerosis is the leading cause of cardiovascular disease, with related events responsible for one-third of all deaths annually^6^. Atherosclerosis is when plaque builds up in the arteries, usually caused by an excess of low-density lipoproteins (LDLs) in the bloodstream. Monocytes are signaled to enter the artery and become macrophages to perform lipid engulfment. When they engulf an excess amount of lipids, macrophages are classified as macrophage-derived foam cells. The buildup of foam cells and plaque restricts blood flow around the body. Artery wall ruptures release plaque into the bloodstream and can block blood flow to critical organs like the brain and heart, which increases the risk of cardiovascular events such as strokes and heart attacks.

The current front-line treatment for preventing atherosclerosis-related cardiovascular events is statin^7^, which lowers cholesterol levels in the bloodstream. Statin and drugs that effectively decrease blood coagulation and hypertension decrease the risk of cardiovascular events but do not eliminate them entirely^8,9^. Modifiable risk factors underscored by inflammation, such as smoking, obesity, and diabetes mellitus, are also targeted in treatment^10,11^. Thus, it is necessary to consider other risk factors in atherosclerosis treatment.

Cardiovascular disease is known to have sexual dimorphism^13^. Men die from atherosclerosis more frequently and develop atherosclerosis at a younger age than women^12^. Meanwhile, women tend to develop atherosclerosis at an older age, particularly during and after menopause transition^14^. In addition, studies have suggested that sex hormones change atherosclerotic immune response to influence disease phenotype^12^. Despite established knowledge of the existence of sex differences in atherosclerosis, there is limited data that thoroughly examines the mechanisms and manifestations of this sex difference^11^. Most existing animal atherosclerosis models either do not consider sex or do not suggest sex differences in animal atherosclerosis^1,12^. Specific knowledge of the impact of sex differences in atherosclerosis development is critical to more effective diagnosis and treatment of atherosclerosis.

Another factor known to affect atherogenesis, but not yet implemented in treatment due to a lack of definitive targets . Several animal and human studies have suggested inflammation as a key factor in atherosclerosis development^15,16,17,18^. For example, the human trial CANTOS showed that targeting inflammatory pathways with canakinumab at a dose of 150 mg every 3 months led to a significantly lower rate of recurrent cardiovascular events than the placebo^15^. Despite the established importance of inflammation in atherosclerosis, atherosclerotic treatments targeting inflammation are uncommon while common therapies mainly target traditional risk factors like lipid-handling dysfunction^19^. Inflammation in atherosclerosis was examined by conducting analysis with whole blood transcriptomics, which included cells that are the precursor of macrophage called monocytes. This paper aims to define sex differences in atherosclerotic inflammation and lipid-handling dysfunction using cross-cohort comparisons of male and female atherosclerosis patients with varying degrees of atherosclerotic severity.

## Methods

### Dataset Background Information

GSE221615 is a dataset available on NCBI’s Gene Expression Omnibus (GEO) Database^20^. Whole blood samples were collected from 391 participants of the Progression of Early Subclinical Atherosclerosis (PESA) study to examine epigenetic age and its correlation with subclinical atherosclerosis (SA). Between June 2010 and February 2014, 4,184 healthy employees of Santander Bank between 40 and 54 years old enrolled in the PESA study. All participants were assessed for either plaque formation in five anatomical territories using 2D/3D vascular ultrasound or for coronary artery calcification (Agatston score ≥ 0.5). The participants were categorized based on the number of sites that contained plaque and coronary artery calcification. The four categories were: No SA, focal SA (one site affected), intermediate SA (two to three sites), or generalized SA (four to six sites). Multi-territory ultrasounds were performed three years after the initial screening to measure plaque and calcification progression. 480 PESA participants, 240 without SA and 240 with SA, were selected for omics analysis based on traditional CV risk factors, age, and sex. The 391 participant samples fulfilled all quality control requirements and were used to obtain whole blood methylomics, transcriptomics, and plasma proteomics. Transcriptomic array intensities were filtered and normalized using the rnb.run.analysis function from RnBeads version 2.0.1(20). Around 800,000 probes were extracted after filtering and then filtered further by the Dasen method.

### Bioinformatics Packages Used for Analysis

Data analysis was conducted in R Studio version 4.4.1 using multiple R packages. dplyr^22^ version 1.1.4, tibble^23^ version 3.2.1, and reshape2^24^ version 1.4.4 packages were used for data manipulation. ggplot2^25^ version 3.5.1 was used for graph generation. DESeq2^26^ version 1.44.0 was used for finding differentially expressed genes.

### Differentially Expressed Gene Collection

Cross-cohort comparisons were done in R Studio using GSE221615’s provided normalized counts, FPKM, and metadata datasets. The FPKM dataset contained a total of 39376 genes from 388 patient samples and was sorted into cohorts based on sex and disease severity(The “Ill” category contained all patient samples that were not “No”). (Table 1) 3 sample IDs were missing from the FPKM matrix dataset and thus were not used, resulting in 388 participants used for further analysis (27 sample IDs for females and 361 sample IDs for males). Fold change was calculated and log2 was transformed for each cohort from the FPKM dataset and was then filtered for row sums above 0, leaving 37502 genes for IPA analysis.

**Table 1.** The number of patient samples in cohorts based on sex and atherosclerotic severity.

|  | Male Participant (n) | Female participant (n) |
| --- | --- | --- |
| Total Samples | 361 | 27 |
| SA Status |  |  |
| No SA | 122 | 14 |
| Focal SA | 48 | 0 |
| Intermediate SA | 54 | 5 |
| Generalized SA | 137 | 8 |
| III SA | 239 | 13 |

### Ingenuity Canonical Pathway Analysis

The filtered fold change dataset was uploaded to QIAGEN Ingenuity Pathway Analysis(IPA). Cross-cohort comparisons were calculated with IPA’s Canonical Pathway. The filtering parameters were a p-value < 0.05 and an absolute value Log 2 ratio ≥ 1.5 (Table 2).

**Table 2.** The number of pathways outputted from IPA’s Canonical Pathway analysis of the filtered FPKM fold change dataset.

|  | Both Sex | Female | Male |
| --- | --- | --- | --- |
| Focal vs Ctrl | 183 | 0 | 183 |
| Intermediate vs Ctrl | 145 | 325 | 347 |
| Generalized vs Ctrl | 133 | 181 | 125 |
| III vs Ctrl | 144 | 192 | 203 |

Female and male outputs were filtered to only contain inflammatory pathways to compare the quantity and enrichment levels of inflammatory pathways between the two sexes(Table 3).

**Table 3:**
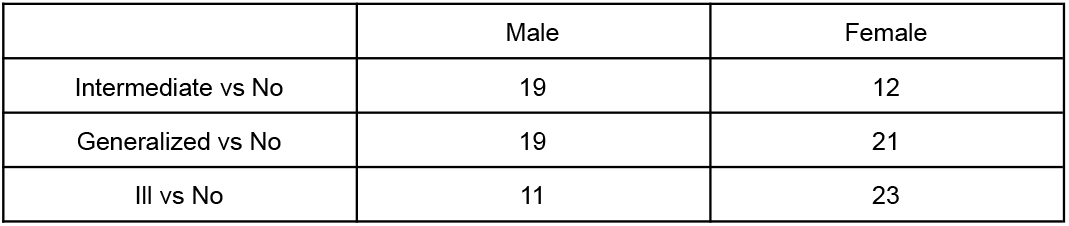
The number of inflammatory pathways in each cohort from Table 2.

|  | Male | Female |
| --- | --- | --- |
| Intermediate vs No | 19 | 12 |
| Generalized vs No | 19 | 21 |
| III vs No | 11 | 23 |

The resulting datasets from males and females were combined by shared cohort comparison and used to create stacked bar charts using ggplot2 (Figure 1A and 1B).

**Figure 1.**
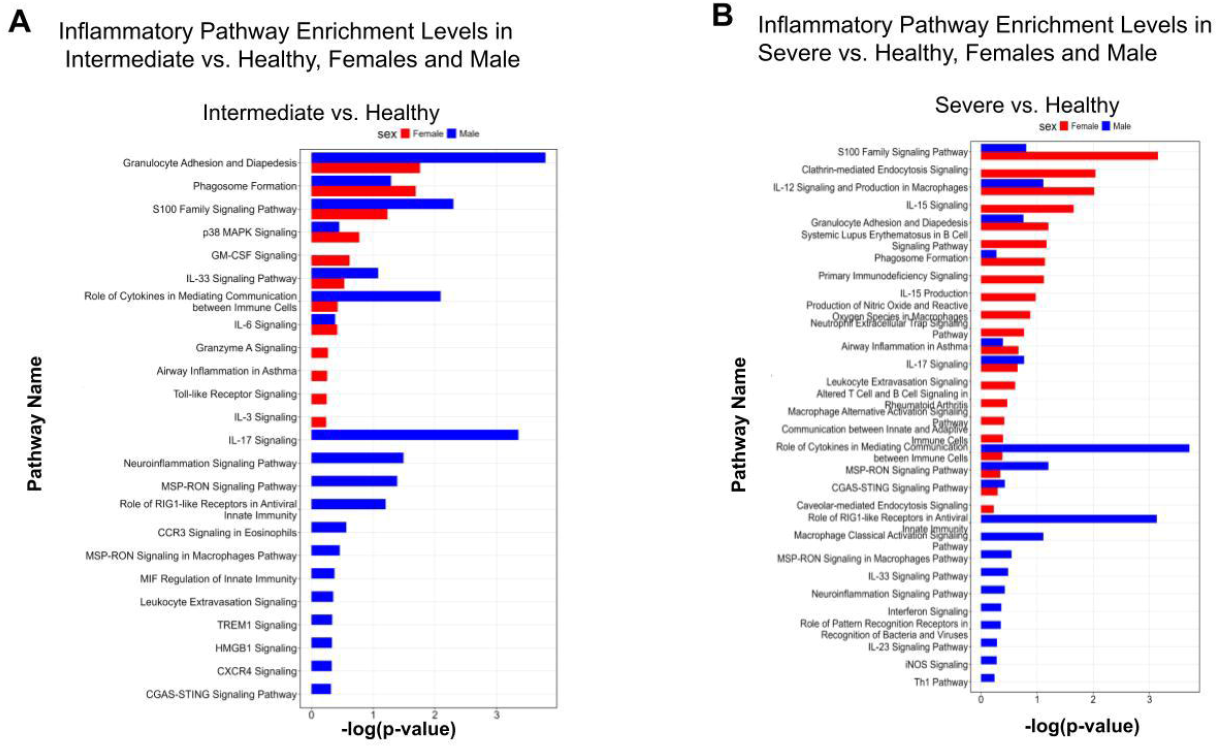
Inflammatory Pathway Signaling is Increased in Females Than Males as Disease Severity Increases. **(A)** Gene ontology analysis using QIAGEN’s Ingenuity Pathway Analysis(IPA) was calculated using GSE221615’s normalized counts FPKM dataset. IPA revealed that in the Intermediate atherosclerosis severity stage, inflammatory pathways are more enriched in males than females. **(B)** Gene ontology analysis using QIAGEN’s Ingenuity Pathway Analysis (IPA) was calculated using GSE221615’s normalized counts FPKM dataset. IPA revealed that in the Generalized/Severe atherosclerosis stage, females have more enriched inflammatory pathways, such as S100 Family Signaling Pathway and IL-12 Signaling and Production in Macrophages, compared to males. This suggests heightened disease severity causes more enriched inflammatory pathways in females.

### AtheroSpectrum

AtheroSpectrum, a single-sample macrophage foaming analysis tool, was used to examine lipid-handling dysfunction and inflammation. The normalized counts dataset was loaded into R for AtheroSpectrum analysis. For each sample, AtheroSpectrum calculated a macrophage polarization index (MPI) value, indicating inflammation level, and a macrophage-derived foam cell index (MDFI) value, indicating lipid-handling level. Samples were categorized based on sex and degree of atherosclerotic severity (PESA score). To compare inflammation and lipid-handling levels between male and female patients across degrees of atherosclerotic severity, MPI and MDFI density plots were generated and 2D density plots were generated based on MPI and MDFI values. Violin dot box plots were generated to compare MDFI levels between female and male patients. Wilcox tests were conducted to determine p-values for MPI and MDFI comparisons between cohorts.

## Results

DEGs were sex-specific across degrees of atherosclerosis, suggesting a difference in male and female atherosclerosis.

### IPA’s Canonical Pathways showed that the role of inflammation in atherogenesis is sex-specific

Data from IPA’s Canonical Pathway output showed that inflammatory pathways were more enriched in females than males as disease severity rose from “Intermediate” to “Generalized” (Figure 1A and 1B). In the Intermediate vs. Healthy comparison, males had more inflammatory and significantly enriched pathways than females (Figure 1A). However, in the Severe vs.

Healthy comparison, inflammatory pathways for females were increased in number (Figure 1B). Some shared pathways that were more expressed in males in the Intermediate vs. Healthy comparison became more expressed in females in the Severe vs. Healthy comparison (S100 Family Signaling Pathway, Granulocyte Adhesion and Diapedesis).

### In line with IPA output, AtheroSpectrum revealed sex differences in atherosclerotic inflammation and lipid-handling dysfunction

AtheroSpectrum analyses revealed that inflammation levels, quantified by MPI, were higher for female patients than male patients with any degrees of atherosclerotic severity (p=0.0223), especially in patients with severe symptoms (p=0.0216) (Figure 2F). In addition, the difference in MPI value between symptomatic female patients and healthy females was greater than the difference in MPI between symptomatic male patients and healthy males across all degrees of atherosclerotic severity, suggesting that inflammation plays a greater role in female atherogenesis than male atherogenesis (Figure 2C and 2D). AtheroSpectrum 2D density plots of MPI and MDFI values revealed that the difference in MDFI between symptomatic male patients and healthy males was greater than the difference in MDFI between symptomatic female patients and healthy females across degrees of atherosclerotic severity (Figure 2A and 2B). Violin dot box plots showing MDFI levels for severe patients versus healthy patients in females and males revealed that MDFI levels were higher for severe male patients than severe female patients, indicating higher lipid-handling dysfunction in male atherosclerosis patients (Figure 2G). Moreover, severe male patients had higher MDFI levels than healthy males, indicating higher lipid-handling dysfunction (p=0.290) with a greater difference than between severe female patients and healthy females (Figure 2G). Thus, AtheroSpectrum results confirmed that inflammation is more important to female atherogenesis than male atherogenesis and showed that lipid-handling dysfunction is more important to male atherogenesis than female atherogenesis.

**Figure 2.**
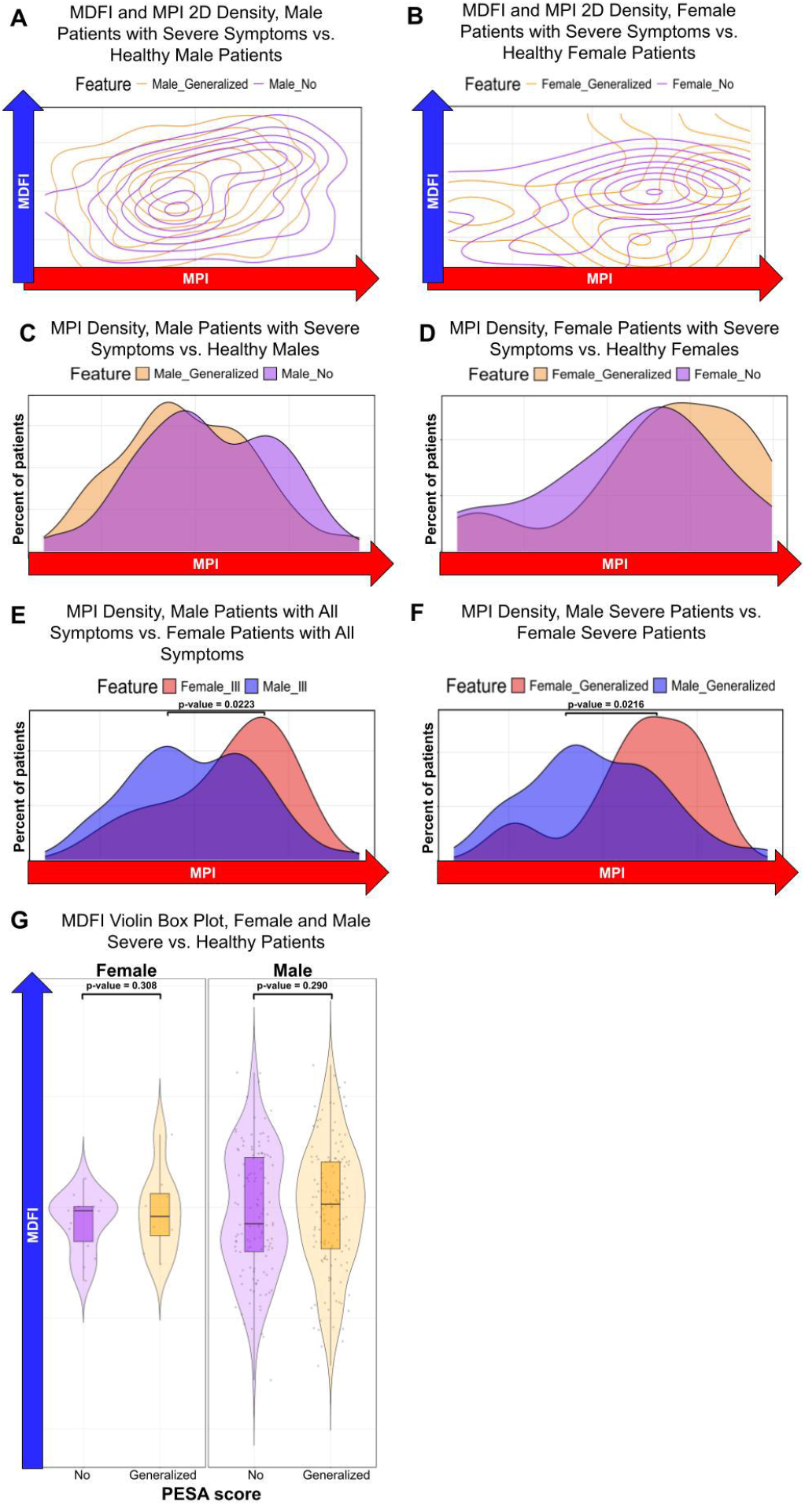
AtheroSpectrum revealed differences in inflammation, quantified by Macrophage Polarization Index (MPI), and lipid-processing, quantified by Macrophage-derived Foaming Index (MDFI), between female and male patients. Normalized counts were inputted into AtheroSpectrum for analysis. **(A)** 2D density MDFI and MPI plots for severe males versus healthy males and **(B)** severe females versus healthy females. Compared to severe females and healthy females, MPI difference between severe males and healthy males is greater, indicating a greater role of lipid-handling in male disease development. (C) MPI density plot of males with all symptoms versus healthy males. **(D)** MPI density plot of females with all symptoms versus healthy females. Compared to males with all symptoms and healthy males, MPI difference between females with all symptoms and healthy females is greater, indicating a greater role of inflammation in female disease development. **(E)** MPI density plot of males with all symptoms versus females with all symptoms. Compared to male patients with all symptoms, female patients with all symptoms had to have higher MPI levels, indicating higher inflammation (p==0.0223) **(F)** MPI density plot of severe males versus severe females. Compared to severe male patients, severe female patients had to have higher MPI levels, indicating higher inf ammation (p==0.0216). **(G)** Violin plots showing MDFI levels for healthy patients and patients with severe symptoms in females and males. Compared to severe female patients, severe male patients had higher MDFI levels, indicating higher lipid-handling dysfunction. Compared to healthy males, severe male patients had higher MDFI levels, indicating higher lipid-handling dysfunction (p==0.290).

Inflammatory pathway activation increased overall sexes as atherosclerotic disease conditions became more severe. However, the level of inflammation differed between male and female atherogenesis. Inflammation in female atherogenesis was more enriched than in male atherogenesis. Conversely, lipid-handling dysfunction levels were higher in male atherogenesis than in female atherogenesis.

## Discussion

Our results confirmed the importance of inflammation in atherosclerosis suggested by several studies, underscoring atherosclerosis as an inflammatory disease rather than just a lipid storage disease as it was previously considered. Thus, inflammation should be targeted in atherosclerosis treatment alongside current targets such as blood coagulation and hypertension. Further, annotated pathways and AtheroSpectrum plots showed differences in the activation of inflammatory pathways in female atherosclerosis patients and male atherosclerosis patients. Inflammation was revealed as more important to female atherogenesis, while lipid-handling dysfunction was more important to male atherogenesis. Thus, while inflammation serves an important role in atherosclerosis across both sexes, patient sex should be a factor considered in the treatment administered to atherosclerosis patients.

The dataset used in our study, GSE221615, has 27 female samples and 361 male samples. The small number of female samples compared to male samples does not reflect the true population sex distribution. Results should be confirmed by replicating our methods using datasets with more female samples or by taking a subset of male samples.

In the future, our results can be used to develop a model to predict atherosclerotic disease and severity in patients based on top DEGs, activation of key pathways, and patient traits such as age and sex. In recent years, the unmet demand for a reliable means of predicting atherosclerotic disease has increased due to recent findings that over forty percent of asymptomatic adults with no known heart disease have undiagnosed atherosclerosis. A successful prediction model would allow a high proportion of adults with subclinical atherosclerosis to receive potentially life-saving treatment early in disease development.

## Acknowledgements

We thank Alyssa J. Matz, a graduate student in the Immunology Department of UConn Health, for her dedication and effort in mentoring us in bioinformatics and paper writing.

## References

1. Man JJ, Beckman JA, Jaffe IZ. Sex as a Biological Variable in Atherosclerosis. Circ Res. 2020 Apr 24;126(9):1297–1319. doi: 10.1161/CIRCRESAHA.120.315930. Epub 2020 Apr 23. PMID: 32324497; PMCID: PMC7185045.

2. AlSiraj, Y., Chen, X., Thatcher, S.E. et al.. XX sex chromosome complement promotes atherosclerosis in mice. Nat Commun 10, 2631 (2019). 10.1038/s41467-019-10462-z

3. Ahn, Y.J., Wang, L., Tavakoli, S. et al.. Glutaredoxin 1 controls monocyte reprogramming during nutrient stress and protects mice against obesity and atherosclerosis in a sex-specific manner. Nat Commun 13, 790 (2022). 10.1038/s41467-022-28433-2

4. Shoaran, M., Maffia, P. Tackling inflammation in atherosclerosis. Nat Rev Cardiol 21, 442 (2024). 10.1038/s41569-024-01007-z

5. Soehnlein, O., Libby, P. Targeting inflammation in atherosclerosis — from experimental insights to the clinic. Nat Rev Drug Discov 20, 589–610 (2021). 10.1038/s41573-021-00198-1

6. Lusis, A. Atherosclerosis. Nature 407, 233–241 (2000). 10.1038/35025203

7. Almeida SO, Budoff M. Effect of statins on atherosclerotic plaque. Trends Cardiovasc Med. 2019 Nov;29(8):451–455. doi: 10.1016/j.tcm.2019.01.001. Epub 2019 Jan 7. PMID: 30642643.

8. Matsuura Y, Kanter JE, Bornfeldt KE. Highlighting Residual Atherosclerotic Cardiovascular Disease Risk. Arterioscler Thromb Vasc Biol. 2019 Jan;39(1):e1–e9. doi: 10.1161/ATVBAHA.118.311999. PMID: 30586334; PMCID: PMC6310032.

9. Wong ND, Zhao Y, Quek RGW, Blumenthal RS, Budoff MJ, Cushman M, Garg P, Sandfort V, Tsai M, Lopez JAG. Residual atherosclerotic cardiovascular disease risk in statin-treated adults: The Multi-Ethnic Study of Atherosclerosis. J Clin Lipidol. 2017 Sep-Oct;11(5):1223–1233. doi: 10.1016/j.jacl.2017.06.015. Epub 2017 Jun 30. PMID: 28754224; PMCID: PMC6394854.

10. Engelen, S.E., Robinson, A.J.B., Zurke, YX. et al.. Therapeutic strategies targeting inflammation and immunity in atherosclerosis: how to proceed?. Nat Rev Cardiol 19, 522–542 (2022). 10.1038/s41569-021-00668-4

11. Herrington W, Lacey B, Sherliker P, Armitage J, Lewington S. Epidemiology of Atherosclerosis and the Potential to Reduce the Global Burden of Atherothrombotic Disease. Circ Res. 2016 Feb 19;118(4):535–46. doi: 10.1161/CIRCRESAHA.115.307611. PMID: 26892956.

12. Fairweather D. Sex differences in inflammation during atherosclerosis. Clin Med Insights Cardiol. 2015 Apr 19;8(Suppl 3):49–59. doi: 10.4137/CMC.S17068. PMID: 25983559; PMCID: PMC4405090.

13. van Dam-Nolen DHK, van Egmond NCM, Koudstaal PJ, van der Lugt A, Bos D. Sex Differences in Carotid Atherosclerosis: A Systematic Review and Meta-Analysis. Stroke. 2023 Feb;54(2):315–326. doi: 10.1161/STROKEAHA.122.041046. Epub 2022 Nov 29. PMID: 36444718; PMCID: PMC9855762.

14. El Khoudary SR, Aggarwal B, Beckie TM, Hodis HN, Johnson AE, Langer RD, Limacher MC, Manson JE, Stefanick ML, Allison MA; American Heart Association Prevention Science Committee of the Council on Epidemiology and Prevention; and Council on Cardiovascular and Stroke Nursing. Menopause Transition and Cardiovascular Disease Risk: Implications for Timing of Early Prevention: A Scientific Statement From the American Heart Association. Circulation. 2020 Dec 22;142(25):e506–e532. doi: 10.1161/CIR.0000000000000912. Epub 2020 Nov 30. PMID: 33251828.

15. Ridker PM, Everett BM, Thuren T, MacFadyen JG, Chang WH, Ballantyne C, Fonseca F, Nicolau J, Koenig W, Anker SD, Kastelein JJP, Cornel JH, Pais P, Pella D, Genest J, Cifkova R, Lorenzatti A, Forster T, Kobalava Z, Vida-Simiti L, Flather M, Shimokawa H, Ogawa H, Dellborg M, Rossi PRF, Troquay RPT, Libby P, Glynn RJ; CANTOS Trial Group. Antiinflammatory Therapy with Canakinumab for Atherosclerotic Disease. N Engl J Med. 2017 Sep 21;377(12):1119–1131. doi: 10.1056/NEJMoa1707914. Epub 2017 Aug 27. PMID: 28845751.

16. Gisterå A, Ketelhuth DFJ, Malin SG, Hansson GK. Animal Models of Atherosclerosis-Supportive Notes and Tricks of the Trade. Circ Res. 2022 Jun 10;130(12):1869–1887. doi: 10.1161/CIRCRESAHA.122.320263. Epub 2022 Jun 9. PMID: 35679358.

17. von Scheidt M, Zhao Y, Kurt Z, Pan C, Zeng L, Yang X, Schunkert H, Lusis AJ. Applications and Limitations of Mouse Models for Understanding Human Atherosclerosis. Cell Metab. 2017 Feb 7;25(2):248–261. doi: 10.1016/j.cmet.2016.11.001. Epub 2016 Dec 1. PMID: 27916529; PMCID: PMC5484632.

18. Hansson GK. Inflammation and Atherosclerosis: The End of a Controversy. Circulation. 2017 Nov 14;136(20):1875–1877. doi: 10.1161/CIRCULATIONAHA.117.030484. Epub 2017 Sep 15. PMID: 28916641.

19. Engelen, S.E., Robinson, A.J.B., Zurke, YX. et al.. Therapeutic strategies targeting inflammation and immunity in atherosclerosis: how to proceed?. Nat Rev Cardiol 19, 522–542 (2022). 10.1038/s41569-021-00668-4

20. Sánchez-Cabo F, Fuster V, Silla-Castro JC, González G, Lorenzo-Vivas E, Alvarez R, Callejas S, Benguría A, Gil E, Núñez E, Oliva B, Mendiguren JM, Cortes-Canteli M, Bueno H, Andrés V, Ordovás JM, Fernández-Friera L, Quesada AJ, Garcia JM, Rossello X, Vázquez J, Dopazo A, Fernández-Ortiz A, Ibáñez B, Fuster JJ, Lara-Pezzi E. Subclinical atherosclerosis and accelerated epigenetic age mediated by inflammation: a multi-omics study. Eur Heart J. 2023 Aug 1;44(29):2698–2709. doi: 10.1093/eurheartj/ehad361. PMID: 37339167; PMCID: PMC10393076.

21. Assenov, Y., Müller, F., Lutsik, P. et al.. Comprehensive analysis of DNA methylation data with RnBeads. Nat Methods 11, 1138–1140 (2014). 10.1038/nmeth.3115

22. Wickham H, François R, Henry L, Müller K, Vaughan D (2023). dplyr: A Grammar of Data Manipulation. R package version 1.1.4, https://github.com/tidyverse/dplyr, https://dplyr.tidyverse.org.

23. Müller K, Wickham H (2023). tibble: Simple Data Frames. https://tibble.tidyverse.org/, https://github.com/tidyverse/tibble.

24. Wickham, H. (2007). Reshaping Data with the reshape Package. In Journal of Statistical Software (Vol. 21, Issue 12, pp. 1–20). http://www.jstatsoft.org/v21/i12/

25. Wickham H (2016). ggplot2: Elegant Graphics for Data Analysis. Springer-Verlag New York. ISBN 978-3-319-24277-4, https://ggplot2.tidyverse.org.

26. Love MI, Huber W, Anders S (2014). “Moderated estimation of fold change and dispersion for RNA-seq data with DESeq2.” Genome Biology, 15, 550. doi:10.1186/s13059-014-0550-8.

